# Functionally non-redundant paralogs *spe-47* and *spe-50* encode FB-MO associated proteins and interact with *him-8*

**DOI:** 10.1101/2020.03.13.990366

**Authors:** Jessica N. Clark, Gaurav Prajapati, Fermina Aldaco, Thomas J. Sokolich, Steven Keung, Sarojani Austin, Ángel A. Valdés, Craig W. LaMunyon

**Affiliations:** Department of Biological Sciences, Cal Poly Pomona, Pomona, California, United States of America

## Abstract

The activation of *C. elegans* spermatids to crawling spermatozoa is affected by a number of genes including *spe-47.* Here, we investigate a paralog to *spe-47*: *spe-50*, which has a highly conserved sequence and expression, but which is not functionally redundant to *spe-47.* Phylogenetic analysis indicates that the duplication event that produced the paralogs occurred prior to the radiation of the *Caenorhabditis* species included in the analysis, allowing a long period for the paralogs to diverge in function. Furthermore, we observed that knockout mutations in both genes, either alone or together, have little effect on sperm function. However, hermaphrodites harboring both knockout mutations combined with a third mutation in the *him-8* gene are nearly self-sterile due to a sperm defect, even though they have numerous apparently normal sperm within their spermathecae. We suggest that the sperm in these triple mutants are defective in fusing with oocytes, and that the effect of the *him-8* mutation is due to its role in chromatin remodeling.

## Introduction

Sperm cells generally face a brief life of intense competition to realize their goal of fertilizing an oocyte. To have success, they must execute with extreme efficiency. They must activate at precisely the right moment, locomote with haste using chemotaxis to guide them to the fertilization site, and fuse with an oocyte as quickly as possible. All this is required of a cell stripped of its ability to express its genome, in most cases surviving only on the meager stores within its tiny volume. Given the unusual nature of sperm cells, it is not surprising that well in excess of 1,000 genes are specific to, or upregulated in, sperm development [1, 2].

Our studies are concerned with the activation of sperm from the nematode *C. elegans.* Spherical and immotile, *C. elegans* spermatids are so primed to activate that they require the activity of SPE-6 to remain in the spermatid stage [3]. Once a signal is received, the spermatids undergo rapid wholesale cellular reorganization that involves an influx of cations [4], a brief elevation in pH [5], the release of intracellular Ca2+ [6-8], induction of a MAPK cascade [9], and polymerization of major sperm protein (MSP) and fusion of the membranous organelles (MOs) with the plasma membrane [7]. As a result, a pseudopod is extended and motility is achieved through MSP mediated pseudopodal treadmilling [10].

As the first step in the life of a *C. elegans* sperm cell, activation (spermiogenesis) may be initiated via two redundant pathways. One pathway, utilized only in males, involves the extracellular signaling serine protease TRY-5, which is secreted with the seminal fluid [11] and activates the spermatid. TRY-5 interacts with the transporter protein SNF-10 to stimulate activation [12]. The second pathway present in both males and hermaphrodites proceeds through the SPE-8 group proteins, namely, SPE-8, SPE-12, SPE-19, SPE-27, SPE-29 [reviewed in 13], and the most recent addition, SPE-43 [14]. It is thought that these proteins are anchored to the plasma membrane and transduce the activation signal inward, perhaps through the non-receptor tyrosine kinase SPE-8, which appears to move inward, away from the plasma membrane, during activation [15].

Our focus has been on discovering the identities of a collection of mutations recovered from a suppressor screen of *spe-27(it132ts)* [3]. Mutant *spe-27* hermaphrodites are sterile because their self-sperm do not activate. The suppressor mutations restore varying degrees of fertility due to the fact that they cause sperm to activate prematurely without the need for activation signal transduction. We have identified *spe-27(it132ts)* suppressor mutations in *spe-4(hc196)* [16], *spe-46(hc197)* [17], and *spe-47(hc198)* [18]. There is a paralog to *spe-47* in the *C. elegans* genome with the sequence identifier Y48B6A.5. Here, we report that this paralog is a new sperm gene designated *spe-50*, but it is not functionally redundant to *spe-47*. However, knockouts of the two genes have an unusual genetic interaction with *him-8* when combined in a triple mutant strain.

## Methods

### Worm strains and handling

All *C. elegans* strains were maintained on *Escherichia coli* OP50-seeded Nematode Growth Media (NGM) agar plates [19]. The *Caenorhabditis* Genetic Center kindly provided the following strains: N2, BA963: *spe-27(it132ts) IV*, BA966: *spe-27(it132ts) unc-22(e66) IV*, CB1489: *him-8(e1489) IV*, DR466: *him-5(e1490) V*, BA17: *fem-1(hc17ts) IV*, JK654: *fem-3(q23ts) IV*, EG5767: *qqIr7 I; oxSi78 II* ; *unc-119(ed3) III*, and SP444 *unc-4(e120) spe-7(mn252)*/*mnC1* [*dpy-10(e128) unc-52(e444)*] *II*. Strain IE4488 harboring the *ttTi4488* Mos1 transposon insertion in Y48B6A.5 was received from the NEMAGENETAG consortium [20], and the transposon insertion was homozygosed to create strain ZQ117. Steven L’Hernault kindly provided BA771 *spe-18(hc133)/mnC1* [*dpy-10(e128) unc-52(e444)*] *II.* Other strains were created by combining alleles. Brood size was measured by counting the progeny laid daily by hermaphrodites isolated in 35 mm petri dishes. In some cases, the effect of mating on hermaphrodite fertility was assessed, in which case individual hermaphrodites were maintained with four males each.

### RT-PCR

To perform RT-PCR, RNA was extracted from mixed-age populations of worms. Large populations of each strain were collected and rinsed 4 times with M9 buffer. After freezing at -80 °C, the worms were disrupted by sonication in TRI reagent, and the RNA was extracted using the Direct-zol™ RNA purification kit following the manufacturer’s protocol for DNase I digestion (Zymo Research). cDNA was synthesized with Maxima Reverse Transcriptase (Thermo Scientific™) and oligo(dT)18 primer (#SO131 Thermo Scientific™). cDNAs were adjusted to give the same concentration across samples prior to PCR amplification. A 536 bp region of *spe-50* cDNA was amplified from exons 3 and 4 with primers that flank the *ttTi4488* Mos1 insertion site (Forward primer: 5’-TTGACTTCTGTGCCTCCAGC -3’; Reverse primer: 5’-GGTTCAACAGATTCTTCCTCAAGTGG-

3’). To determine if gene expression was upregulated in sperm, we multiplexed *spe-50* specific primers with primers that amplify an 898 bp region of the transcript of *act-2*, the *C. elegans* ortholog of β-Actin (Forward primer: 5’-GTATGGGACAGAAAGACTCG-3’; Reverse primer: 5’-ATAGATCCTCCGATCCAGAC-3’). The primers spanned intronic regions to distinguish between products from genomic DNA and cDNA. To determine if the *ttTi4488* Mos1 transposon insertion disrupted *spe-50* transcription, we compared RT-PCR from a population of *spe-50(ttTi4488)* with that from an N2 population. To determine if the *spe-50* transcript is upregulated in sperm, we compared RT-PCR products from populations of *fem-3(q23ts)*, hermaphrodites of which make only sperm, with products from *fem-1(hc13ts)*, hermaphrodites of which make only oocytes.

### CRISPR/Cas9-induced mutations

To induce specific mutations, we utilized the co-conversion strategy for CRISPR/Cas9 mediated gene edits [21]. Briefly, this strategy induces the dominant *cn64* mutation in the *dpy-10* gene in addition to the desired gene-specific edit. F1 worms heterozygous for *cn64* roll while they crawl and are more likely to also harbor the desired edit than do non-rollers [21]. Our two specific edits were accomplished by different methods. To create a mutation that replicates the *spe-47(hc198)* amino acid substitution in *spe-50*, we utilized the expression vector pDD162, which has both a single guide RNA (sgRNA) backbone and the Cas9 gene for *C. elegans* expression [22] (obtained from Addgene). The *spe-50* target sequence (5’-GATCTTGTTACAGTTCCAT-3’) was chosen using a CRISPR guide-finding feature in Geneious R11 (https://www.geneious.com) based upon a high predicted on-target activity [23] and a low likelihood of off-target activity. The targeting sequence was inserted into the sgRNA cassette of pDD162 using the Q5® Site-Directed Mutagenesis Kit (New England BioLabs, Inc.), resulting in identical plasmids (pTS11 and pTS12).

To make the specific *spe-50* edit at the Cas9-induced double-stranded break, we designed an asymmetric ssDNA repair oligonucleotide having 34 bases upstream and 60 bases downstream of the beginning of the PAM site [24]. The oligo had a two-base pair (bp) substitution that changed the Asn at position 314 to Ile, a second silent substitution that created a TaqI restriction site, and a third silent substitution that disrupted the PAM site (Fig. 1C). The *dpy-10* co-conversion edit was accomplished with pDD162 derivative plasmids (pTS5 and pTS6) harboring the *dpy-10* guide. The ssDNA repair oligo to induce the *dpy-10(cn64)* dominant mutation was as described in ARRIBERE *et al.* [21]. The *spe-50* and *dpy-10* CRISPR/Cas9 plasmids and repair oligos were injected into the gonads of N2 hermaphrodites.

**Fig 1.**
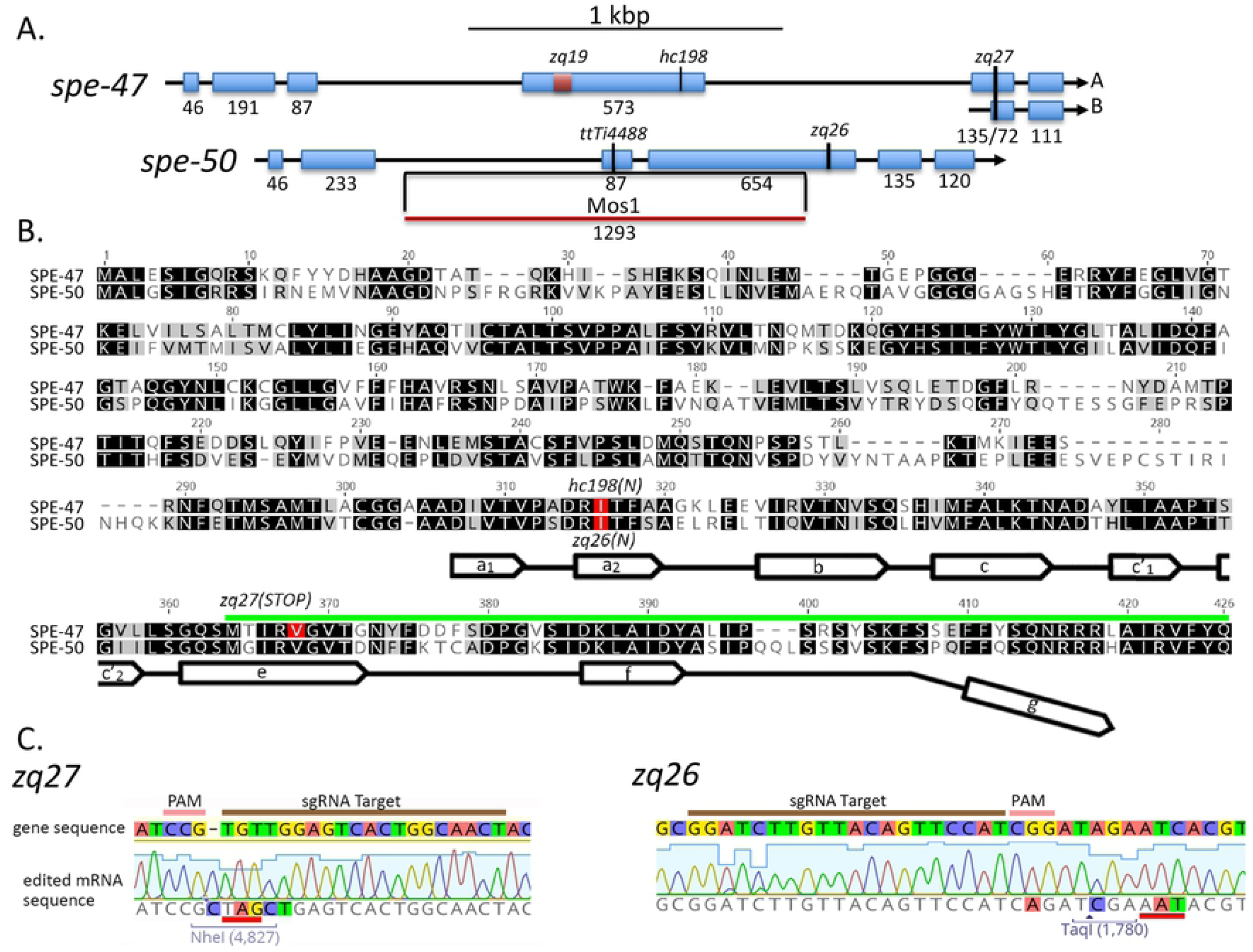
Comparison of *spe-47* and *spe-50* sequences. (A) Exonic structure of *spe-47* and its paralog *spe-50*. Shown are the locations of sequence variants for these genes. (B) Alignment of the SPE-47 and SPE-50 protein sequences. Darker background indicates greater similarity, and mutations are shown in red. Near the carboxy terminus a line with open arrows indicates the partial MSP domain, where the open arrows correspond to the seven β strands present in *A. suum* MSP-α. Note that the MSP domains are truncated at the carboxy terminus, missing the segment indicated by the downward bend in the MSP marker line. Also, the region encoded by the *spe-47* isoform B is indicated by the green line. (C) Two mutations created in this study. The original gene sequence is shown above with the edited mRNA sequence below. The *zq27* mutation created in *spe-47* creates a stop codon (underlined in red) within an NheI restriction enzyme site for detection and a 1 bp insertion to shift the reading frame. The *zq26* mutation in *spe-50* induced an Asn to Ile mutation in the position corresponding the *hc198* mutation in *spe-47*. Sequencing traces shown confirmation that the sequences were edited in the mutant strains.

A knockout mutation that affects both isoforms of *spe-47* was accomplished with the Alt-R™ CRISPR/Cas9 components (Integrated DNA Technologies™) for *in vitro* assembled Cas9-crRNA-tracrRNA ribonucleoproteins (RNPs) following the protocol of KOHLER *et al.* [25]. The *spe-47* target sequence was chosen using the CRISPR guide-finding feature in Geneious R11 (5’-AGTTGCCAGTGACTCCAACA-3’). A specific *spe-47* edit that affects both spliced isoforms was designed into an ssDNA repair oligo. The oligo consisted of 55 bp upstream and 58 bp downstream of the beginning of the PAM site. The oligo had an altered sequence that disrupted five of the six bp just upstream of the PAM site, induced an NheI site that created an in frame stop codon, and inserted one bp to shift the reading frame (Fig. 1).

To induce the *spe-47* knockout, we incubated equimolar solutions of our target-specific crRNAs (for both *spe-47* and *dpy-10*) and standard tracrRNA (100 µM each) in IDT Nuclease-Free Duplex Buffer at 95 °C for 5 minutes followed by 5 minutes at room temperature. The RNA duplex and Cas9-NLS were combined for a final concentration of 27 µM each and incubated at room temperature for 5 minutes to form the final ribonucleoprotein (RNP). We injected worm gonads with a mixture of 17.5 µM RNP and 6 µM ssDNA repair template for *spe-47* along with 0.5 µM ssDNA repair template for *dpy-10*.

To recover the *spe-50* edit, 38 F1 rolling *cn64/+* worms were recovered and isolated. After laying eggs, the F1 worms placed in tubes; their DNA was then extracted and used as template in 5 µl PCR reactions with primers that flank the edit site (Forward primer: 5’-CATCAAGGGTGGACTTCTCG-3’; Reverse primer: 5’-AGCAGCAATGAGATGAGTGTCC-3’). To the completed PCR reactions, we added 5 µl containing restriction enzyme buffer and 5 units of TaqI. After an hour of incubation, the components were run on agarose gels to determine if the edit was induced. Of the 38 F1 rollers isolated, four appeared to have the edit via their restriction digested PCR products. Only one of them was pursued, and it contained the correct alteration of base pairs (Fig 1).

For *spe-47*, after injecting the constituted CRISPR/Cas9 RNPs, we found no F1 rollers. We combined three non-rolling F1s per petri dish in eight dishes, and extracted their combined DNA for PCR/restriction analysis as previously described. One plate appeared to harbor a mutant. After isolating 24 offspring from this plate, we recovered a single worm that was homozygous for the edit (Fig 1). These mutations were designated *spe-50(zq26)* and *spe-47(zq27).*

### Construction of a *spe-50* translational reporter

We created an N-terminal mCherry translational reporter construct for *spe-50* following the MosSCI technique [26, 27]. The mCherry sequence, amplified without its stop codon from plasmid pCFJ104 (Addgene), was placed directly downstream of 1,714 bp of the *spe-50* promoter sequence and was followed by the *spe-50* genomic sequence and 448 bp of the 3’ UTR. All worm sequences were amplified from N2 DNA, and all PCR was performed with Phusion High Fidelity DNA Polymerase (Thermo Scientific). The sequences were amplified with PCR primers engineered with regions of ∼20 bp overlap, enabling us to join them together following the PCR fusion technique described by Hobert [28]. The final fusion was cloned into the multiple cloning site of the vector pCFJ352, which targets the *ttTi4348* Mos1 insertion on Chromosome I for homologous recombination.

### Microscopy, in vitro sperm activation, and microinjection transformation

Imaging was accomplished on a Nikon C2 confocal microscope also outfitted for Nomarski DIC and widefield epifluorescence. Widefield images were captured on a Nikon DS-Qi1 12 bit monochrome camera. Images were acquired and analyzed with Nikon NIS-Elements imaging software. All worms were dissected in SM1 buffer[29], and nuclear material was labeled with 30 ng/µl Hoechst 33342 in SM1 for live cells and with 20 ng/µl DAPI in PBS for fixed and permeabilized cells. Sperm were activated *in vitro* by exposure to SM1 containing 200 µg/ml Pronase. Imaging of reporter constructs was kept constant across experiments to reduce error (e.g. the laser power and gain were used for each fluorophore/fluorescent label). Compounds were microinjected of into the gonads of recipient young adult hermaphrodites using a Nikon Eclipse Ti inverted microscope outfitted for Nomarski DIC.

### Phylogenetic analysis

In order to estimate the evolutionary relationships of the SPE-50 homologous proteins and detect gene duplication events, we conducted a phylogenetic analysis using 15 protein sequences from 7 species of *Caenorhabditis*, with the protein OVOC10046 of *Onchocerca volvulus* as the outgroup. The analysis was run in MrBayes 3.2.6 [30] with the GTR + I model and two runs of six chains for 10 million repetitions, with a sampling interval of 1,000 repetitions and burn-in of 25%.

## Results

### *spe-50* is a sperm gene and overlaps in expression with *spe-47*

When we first discovered that *spe-47* harbored the *hc198* mutation that suppressed *spe-27(it132ts)* sterility by inducing premature spermatid activation [18], we became aware that there was a closely-related paralog present in the genome: Y48B6A.5 (Fig 1 A&B). The SPE-47 and SPE-50 proteins exhibit a high degree of sequence conservation, with both having an N-terminus of unknown function and a C-terminal MSP domain that lacks the final ß strand (Fig 1 B). To determine if Y48B6A.5 expression is upregulated in sperm, we performed differential RT-PCR. The Y48B6A.5 transcript is abundant in *fem-3(q23)* mutant hermaphrodites (Fig 2); these worms produce only spermatids but are otherwise somatically hermaphrodites. Alternatively, the transcript is nearly absent in *fem-1(hc13ts)* hermaphrodites that produce only oocytes. This pattern is characteristic of sperm genes.

**Fig 2.**
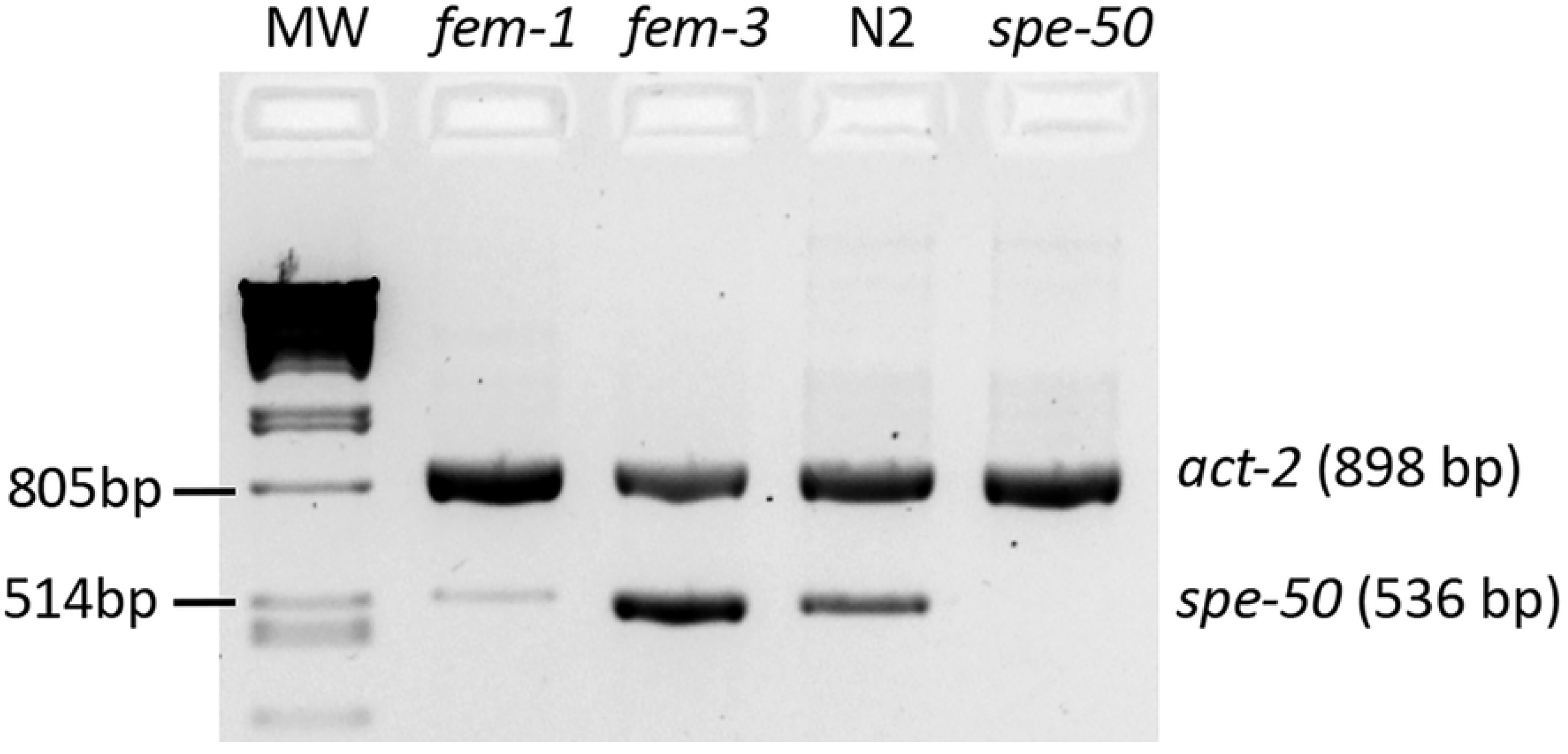
RT-PCR results for the *spe-50* transcript. Primers specific for the Y48B6A.5 transcript amplified a robust product in *fem-3(q23ts)* hermaphrodites that produce only sperm, but such a product was nearly absent amplifying from *fem-1(hc13ts)* hermaphrodites that produce only oocytes. The *spe-50* transcript was also present in the N2 strain but not in the *spe-50(ttTi4488)* mutant that has the Mos1 transposon insertion in Exon 3. Had the transcript with the Mos1 transposon been amplified, it would have been 1,829 bp in length, and the extension time was designed to allow a product that large to be amplified. The PCR reactions also had primers for *act-2*, the *C. elegans* β-actin gene. The *act-2* product demonstrates that there was equivalent mRNA present in the samples. MW is the molecular weight marker: Phage lambda DNA digested with PstI.

There are two sperm genes mapped to the region of Y48B6A.5: *spe-7* [31] and *spe-18* (Steven L’Hernault, personal communication). In order to test whether Y48B6A.5 is actually one of the two nearby genes, we conducted complementation tests using the strain ZQ117 with the *ttTi4488* Mos1 transposon insertion in Y48B6A.5. This insertion disrupts Y48B6A.5 and results in the absence of a transcript (Fig 2). Hermaphrodites homozygous for mutations in *spe-7(mn252)* and *spe-18(hc133)* are sterile due to primary spermatocytes that arrest in Meiosis I [31; Steven L’Hernault, personal communication]. Males from the *ttTi4488* bearing strain were crossed with sterile *unc-4 spe-7* mutant hermaphrodites or with sterile *spe-18* mutant hermaphrodites. The F1 hermaphrodites were isolated at 25 °C and their progeny counted. The F1 hermaphrodites had wild-type fertility: for *spe-7*, F1 fertility = 198 progeny (n=10, SEM=14.4), and for *spe-18*, F1 fertility = 190 progeny (n=12, SEM=9.3). Thus, the *ttTi4488* strain complemented both *spe-7* and *spe-18* because it carried wild-type alleles of both. Y48B6A.5 is a new sperm gene and was given the designation *spe-50* (Steven L’Hernault, personal communication).

To examine SPE-50 protein localization, we created an N-terminal translational reporter with mCherry via the mosSCI protocol [26, 27]. The mosSCI process inserts the reporter into specific chromosomal locations, allowing us to combine the *spe-50::mCherry* reporter with a *spe-47::GFP* reporter we created earlier [18] in a double reporter strain. Imaging of male gonads showed that SPE-50::mCherry colocalizes almost completely with SPE-47::GFP (Fig 3A). Both appear as small puncta surrounding nuclei that are entering the pachytene stage. The puncta enlarge and expand to fill the cells as they mature into primary spermatocytes. Both also then disappear as the secondary spermatocytes form with the spermatids being completely devoid of the reporters. No such fluorescence was found in males lacking the reporters (Fig 3B).

**Fig 3.**
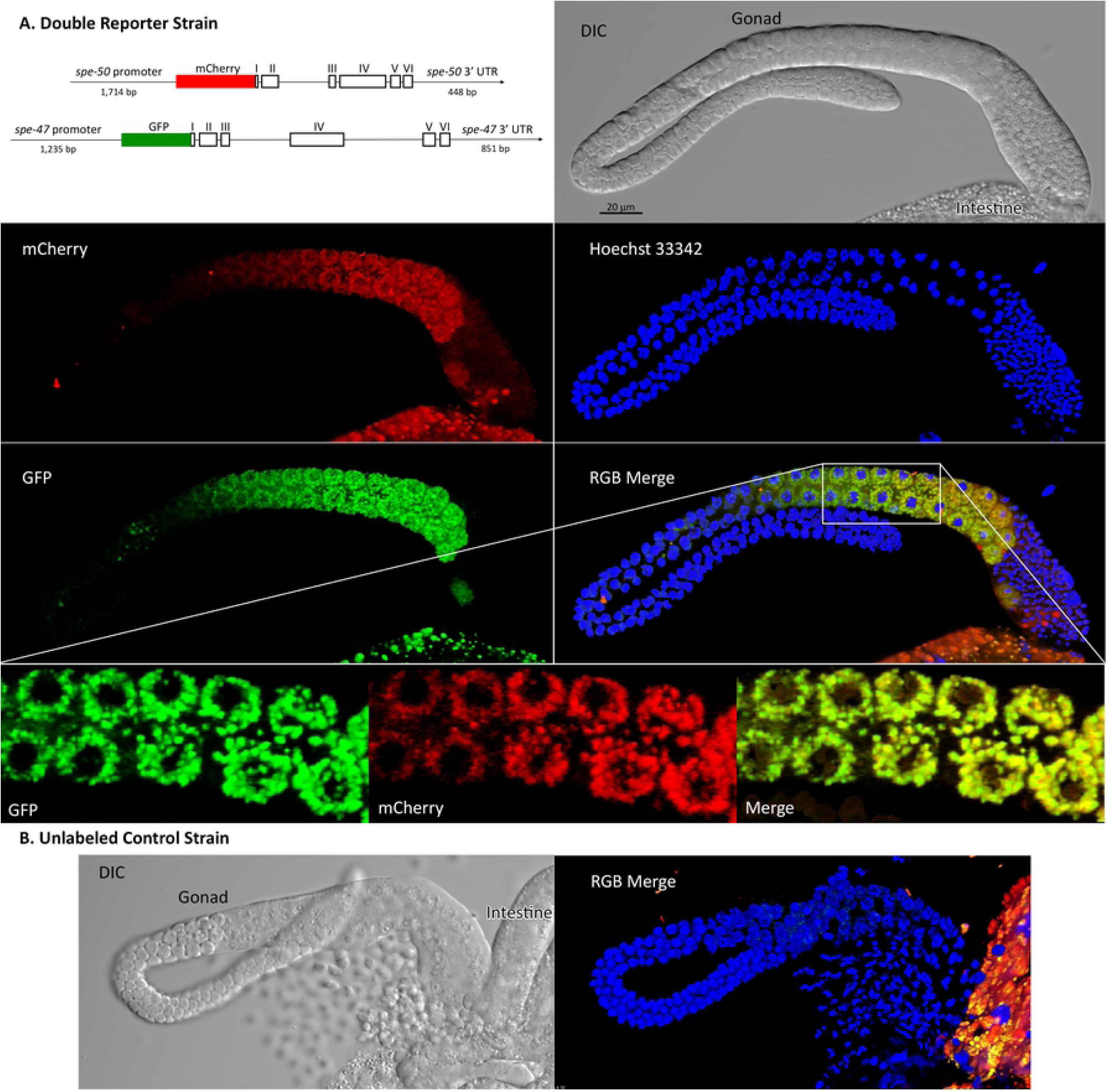
Localization of SPE-50::mCherry and SPE-47::GFP translational reporters in male gonads. The fluorescent images are 3D reconstructions of a stack of images. (A) The double reporter strain constructs and imaging in blue (nuclei), green (*spe-47::GFP*), and red (*spe-50::mCherry*). A region of the gonad in the merge image shown by the box is enlarged to give better detail of the localization. In this enlargement, only the middle of the 3D reconstruction is shown to give better understanding of colocalization. (B) Imaging from the unlabeled wild-type strain for comparison. In both the reporter images and the wild-type control, there are remnants of the intestine present. The intestine is highly autofluorescent in green and red.

The colocalization of the two reporters in space and time suggested that these proteins are involved in the same cellular processes. We tested this hypothesis by examining mutations in both genes. The *spe-47* gene was discovered in a suppressor screen of *spe-27(it132ts)*. The *spe-27* mutation causes hermaphrodite sterility because the self-spermatids are unresponsive to the signal to activate [32]. The *spe-47(hc198)* mutation recovered from the screen causes some spermatids to activate precociously without the need for an activation signal, overcoming the sterility of *spe-27(it132ts)* (Fig 4) [18]. If SPE-50 is functionally redundant to SPE-47, then a mutation similar to *hc198* in *spe-50* should also result in precocious spermatid activation. Fig 1B shows the location of the amino acid changed by *spe-47(hc198)*: an isoleucine to asparagine substitution in the a_2_ ß-strand of the MSP domain. Using CRISPER/Cas9, we induced the same amino acid change in *spe-50* by creating the *zq26* mutation. In addition to altering the two base pairs that cause the amino acid change, we made two other silent substitutions: one that created a TaqI restriction site to allow detection of the alteration, and the other that altered the PAM site to eliminate further Cas9 activity (Fig 1C). Interestingly, the *spe-50(zq26)* mutation did not suppress *spe-27(it132ts)* sterility (Fig 4). In fact, there was no fertility deficit associated with the *spe-50(zq26)* mutation, while *spe-47(hc198)* causes a significant reduction in fertility due to problems with sperm function (Fig 4) [18]. When both *spe-47(hc198)* and *spe-50(zq26)* were combined in the same strain, the fertility was nearly identical to that of *spe-47(hc198)* alone (Fig 4). Thus, in terms of function, the two genes are not identical.

**Fig 4.**
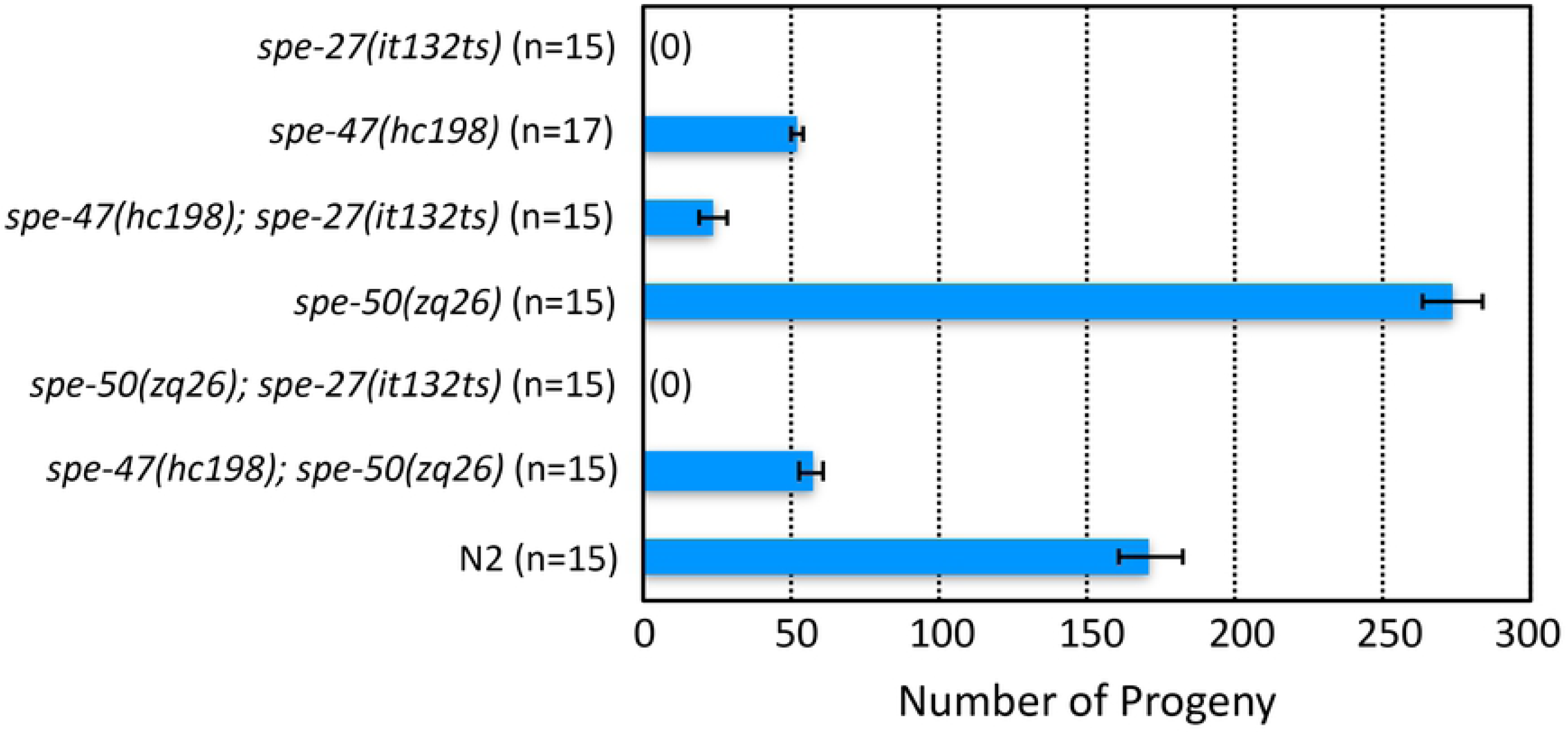
Suppression of *spe-27(it132ts)* sterility. *spe-27(it132ts)* mutants are sterile at 25 °C, but they regain some fertility if they are also homozygous for the *spe-47(hc198)* mutation. On its own, the *hc198* mutation results in a loss of fertility compared to wild type (N2). Conversely, the *spe-50(zq26)* mutation, which encodes the equivalent amino acid change as *hc198*, is entirely fertile on its own and does not suppress *spe-27(it132ts)* sterility. A strain with both *hc198* and *zq26* has approximately the same fertility as *hc198* on its own. Thus, the *spe-50(zq26)* mutation has no apparent effect on fertility.

We also examined knockout alleles of the two genes (Fig 1). Even though the *spe-50(ttTi4488)* transposon insertion disrupts *spe*-*50*, it had essentially no effect on fertility (Fig 5). In our previous study of *spe-47*, we created a knockout allele, *spe-47(zq19)* (Fig 1A), which caused only a slight reduction in fertility [18]. In the interim, a second isoform of the *spe-47* transcript, which is unaffected by the *spe-47(zq19)* mutation, was identified. To ensure that we disabled both isoforms, we inserted a stop codon and frame-shift in the fifth exon common to both isoforms to create a new knockout allele: *spe-47(zq27)* (Fig 1A and 1C). This new mutation also had little effect on fertility (Fig 5). Combining both *spe-50(ttTi4488)* and *spe-47(zq27)* mutations in the same strain of worms had only a modest effect on fertility, reducing it to just over 100 self-progeny per worm (Fig 5). Thus, these genes are not essential to spermatogenesis, but they do seem to have an interaction that reduces fertility when both gene products are absent.

**Fig 5.**
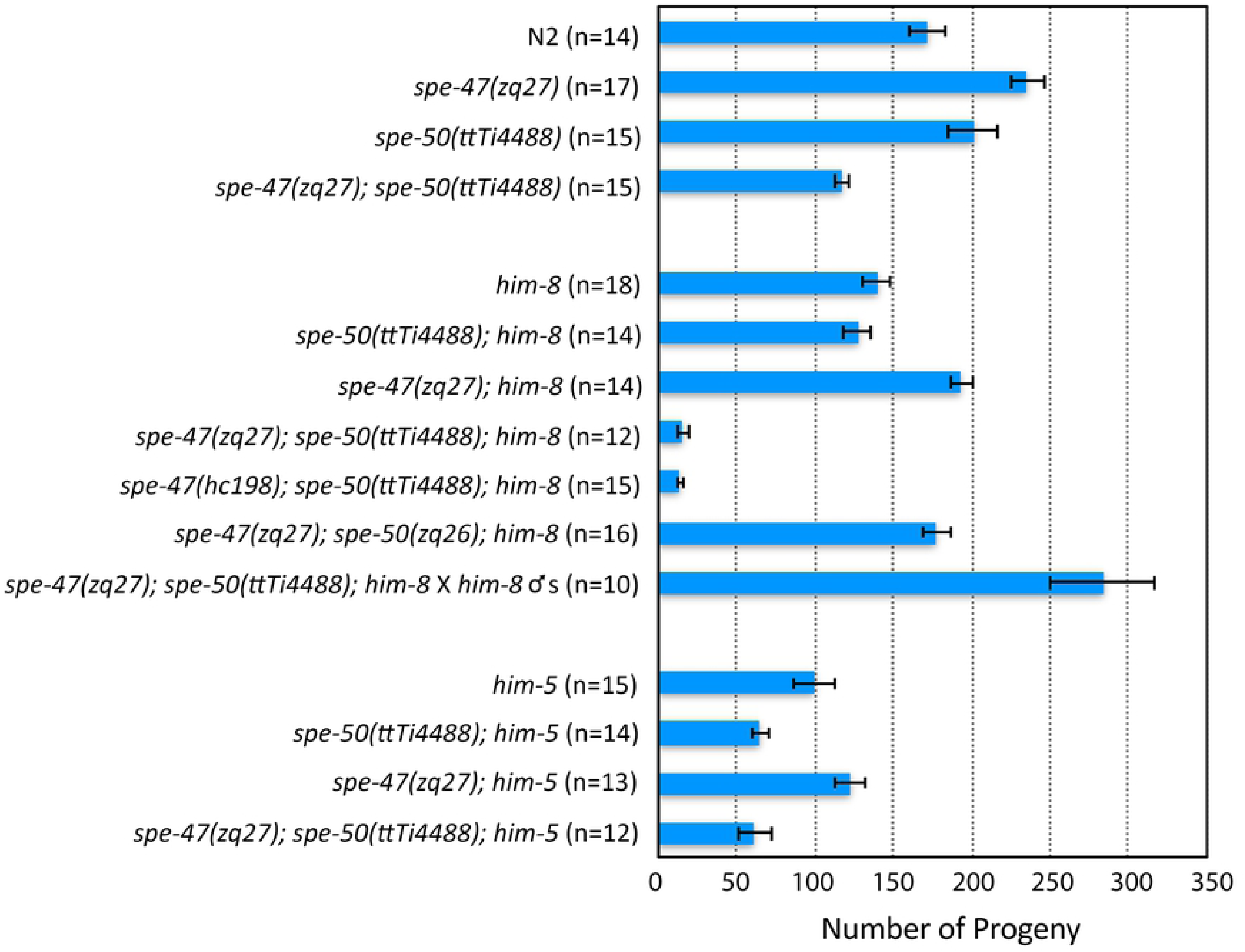
Fertility associated with *spe-47* and *spe-50* knockout mutations and *him* mutant backgrounds. In the top set of bars, the two knockout mutations are compared to N2 both alone and combined in the same strain. In the middle set of bars, the various knockout and *spe-27* suppressor mutations are combined with *him-8(e1489).* Both *spe-47* mutations in a *spe-50* knockout and *him-8* knockout background resulted in a drastic reduction of fertility. This reduction was due to a defect in sperm, because mating the strain to *him-8* males restored full fertility. In the bottom set of bars, combining the knockout mutations with *him-5* did not have a drastic reduction in fertility.

In creating strains, we noticed that combining the *him-8(e1489)* mutation with the two knockout mutations had a more profound effect on fertility (Fig 5). Combining either knockout alone with *him-8(e1489)* did not reduce fertility more than what we found in the *him-8(e1489)* strain alone. It was only when both knockouts were present in the *him-8* background that fertility was reduced approximately 15 progeny per hermaphrodite (Fig 5). This fertility deficit was due to self-sperm dysfunction, as mating these hermaphrodites to *him-8(e1489)* males increased their fertility greatly (Fig 5). Interestingly, combining the *spe-27-*suppressor mutation *spe-47(hc198)* with *spe-50(ttTi4488)* in a *him-8(e1489)* background lead to a similar drop in fertility. The same was not true when we combined the *spe-47(zq27)* knockout mutation with the *spe-27-*suppressor-like mutation *spe-50(zq26)* in a *him-8(e1489)* strain: the fertility was much higher (Fig 5). There was no similar fertility deficit associated with *him-5.* The triple *spe-47(zq27); spe-50(ttTi4488); him-5(e1490)* mutant hermaphrodites laid in excess of 50 offspring (Fig 5), indicating that the interaction with *him-8* is not due just to X Chromosome non-disjunction problems common to both *him-5* and *him-8* mutants.

If the two knockout mutations in a *him-8(e1489)* background (the triple mutant) have defective sperm, then we might expect to see some defects in the sperm themselves. Of 145 sperm dissected from seven triple mutant males, all appeared as normal spermatids (Table I). This is similar to 162 sperm we dissected from five *him-8(e1489)* virgin males: all were spermatids. Because *him-8* has a role in X-Chromosome pairing and synapsis [33], we also looked for gross abnormalities in sperm nuclei. The vast majority of sperm from triple mutants had normal nuclei, similar to what we found for sperm from *him-8(e1489)* single mutants (Table I). Alternatively, spermatids from triple mutants could have defective activation, so we exposed spermatids to the *in vitro* activator Pronase [5, 34]. Sperm from triple mutants kept at 25 °C activated at a slightly but significantly reduced rate (88.5%) compared with sperm from *him-8(e1489)* mutants (96.3%) (Table I), although this does not seem a large enough effect to explain the fertility deficit in this strain. Finally, we looked at the sperm remaining in hermaphrodites one day after being transferred as L4s from 20 °C to 25 °C. The triple mutants had fewer sperm remaining in each gonad arm (mean = 31.7, SEM = 5.9, n = 21 gonad arms) than did *him-8* mutants (mean = 73.5, SEM = 6.9, n = 14 gonad arms), a significant difference (*t*=4.88, *P*<0.001). Again, this does not explain the small number of fertilized eggs produced by the triple mutants, because the number of sperm cells remaining per gonad arm is greater than the number of progeny produced by the triple mutants. In most instances, the sperm in the triple mutants were in or very near the spermatheca, while in others some sperm were well away from the spermatheca, being scattered in the uterus, as if they were unable to remain localized in their target organ. Thus, overall there were more sperm remaining within triple mutants than the number of fertilized eggs they produce, suggesting that these sperm are unable to fertilize oocytes.

We examined the phylogeny of *spe-50* and *spe-47* by comparing their protein products to those from closely related species. The evolutionary analysis in Fig 6 shows that the SPE-50 protein is clearly more closely related to orthologous proteins in the genus than to its paralog-encoded SPE-47. Indeed, the hypothetical duplication of the ancestral coding sequence must have taken place prior to the radiation of the species examined.

**Table I:**
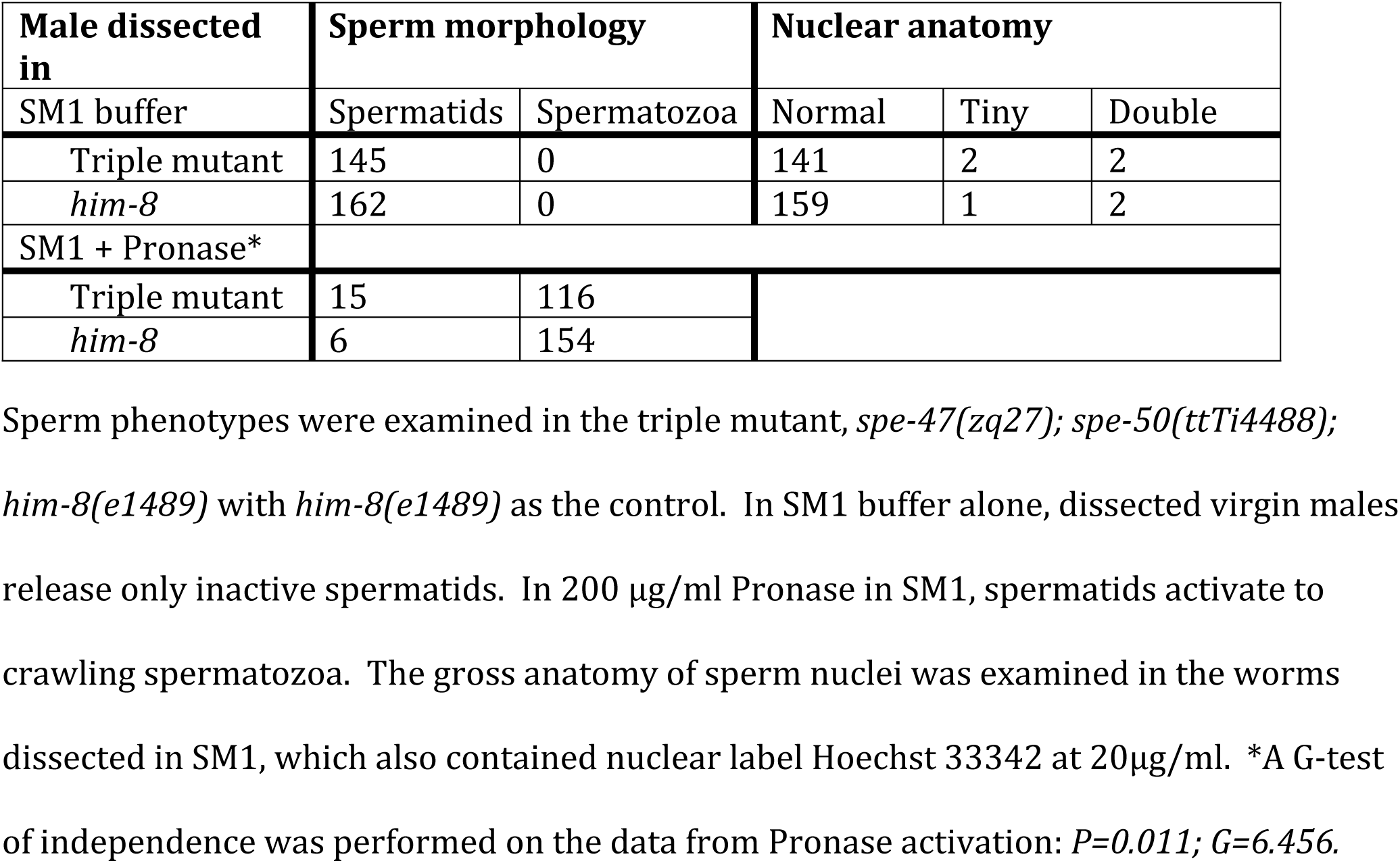
Sperm activation and nuclear anatomy.

| Male dissected in | Sperm morphology |  | Nuclear anatomy |  |  |
| --- | --- | --- | --- | --- | --- |
|  | Spermatids | Spermatozoa | Normal | Tiny | Double |
| SM1 buffer |  |  |  |  |  |
| Triple mutant | 145 | 0 | 141 | 2 | 2 |
| *him-8* | 162 | 0 | 159 | 1 | 2 |
| SM1 + Pronase* |  |  |  |  |  |
| Triple mutant | 15 | 116 |  |  |  |
| *him-8* | 6 | 154 |  |  |  |

Sperm phenotypes were examined in the triple mutant, *spe-47(zq27); spe-50(ttTi4488); him-8(e1489)* with *him-8(e1489)* as the control. In SM1 buffer alone, dissected virgin males release only inactive spermatids. In 200 µg/ml Pronase in SM1, spermatids activate to crawling spermatozoa. The gross anatomy of sperm nuclei was examined in the worms dissected in SM1, which also contained nuclear label Hoechst 33342 at 20µg/ml. \*A G-test of independence was performed on the data from Pronase activation: $P=0.011$ ; $G=6.456$ .

**Fig 6.**
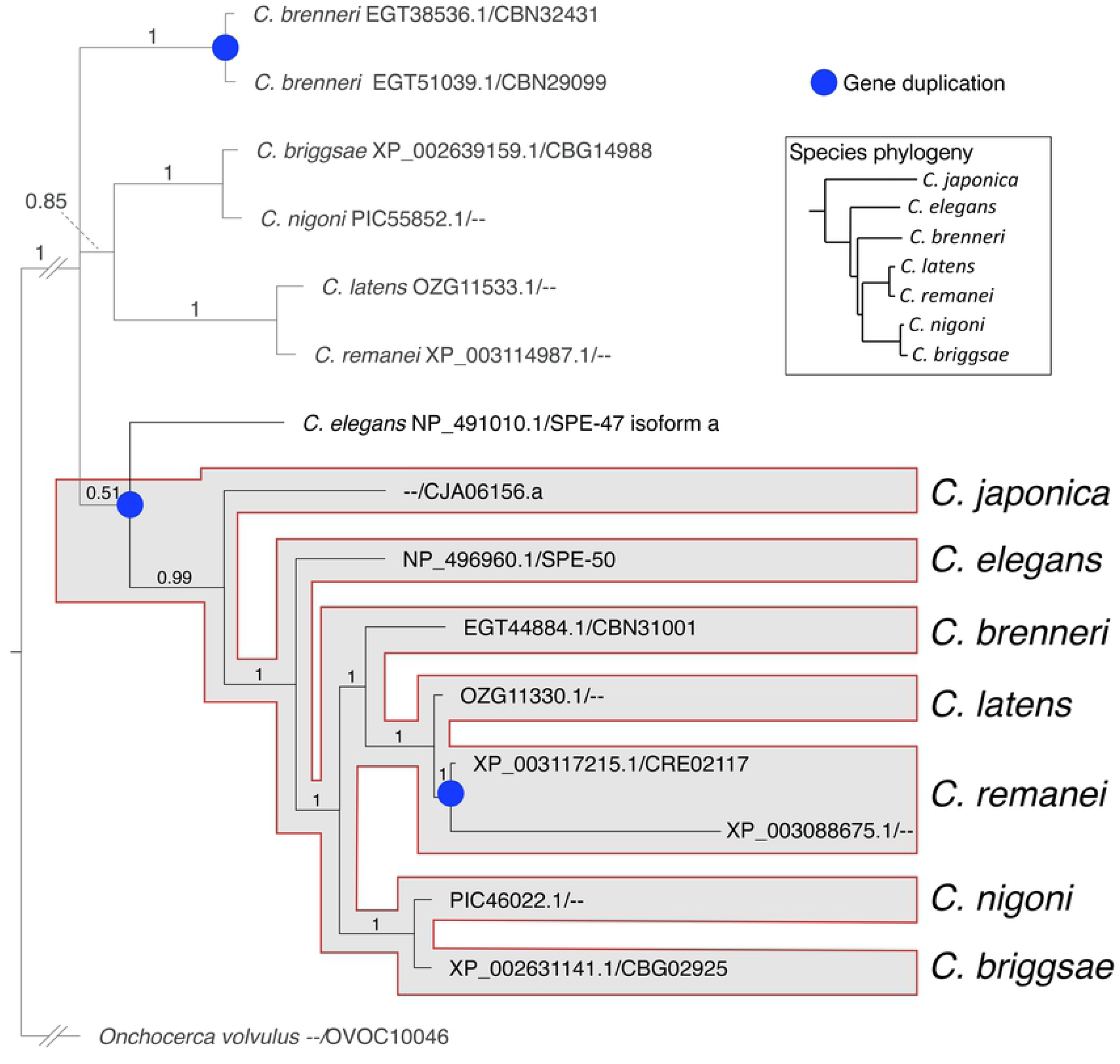
Evolutionary relationship of the SPE-50 homologous proteins. The proteins were identified from a BLASTP search of the NCBI non-redundant protein sequence database. For each protein, the accession number is listed before the slash, and the WormBase identifier after the slash. The duplication event that resulted in creation of the two paralogs occurred prior to the radiation of the species included in the analysis. The phylogeny of the species within *Caenorhabditis* follows STEVENS *et al.* [35].

## Discussion

The paralogs *spe-47* and *spe-50* have a high degree of protein sequence conservation, and they retain a very similar exonic structure (Fig 1). Further, the gene products are expressed in a nearly identical fashion within the spermatogenic tissue. Such similarity would suggest a similar function. However, the two genes are not functionally redundant, as the *spe-50(zq26)* mutation did not phenocopy its homologous *spe-47(hc198)* mutation in suppressing *spe-27(it132ts)* sterility. Also, a strain with knockout mutations in both genes had only slightly reduced fertility. Hypothetically, genes have multiple selection pressures acting on their function, and these pressures may act in opposition, constraining sequence evolution [36]. After a gene duplication event, the paralogs are thought to undergo functional divergence to satisfy different selective pressures with opposing effects on sequence evolution. Thus, true functionally redundant paralogs are very rare [36]. The duplication event that gave rise to *spe-47* and *spe-50* occurred early in the radiation of the genus (Fig 6), so the two paralogs have had ample time to evolve in response to different selection pressures.

However, the *spe-47*and *spe-50* genes do show a phenotype when the knockout mutations are combined in a triple mutant strain with *him-8(e1489).* Fertility in the triple mutant strain is dramatically reduced due to a sperm defect. HIM-8 is a C2H2 zinc-finger protein that binds to the pairing center of the X-chromosome and initiates the pairing and synapsis of the X chromosome homologs [37]; *him-8* mutants show high levels of X-chromosome nondisjunction leading to increased rates of male production. HIM-8 protein binds to specific short sequences concentrated in the pairing centers, but these binding sequences are present at other sites on the X-chromosome and on the autosomes [38]. Thus, it is not surprising that HIM-8 protein is also bound more diffusely at other sites on the X chromosome and the autosomes [39]. Further, mutations in *him-8* can suppress the defects associated with hypomorphic mutations in *egl-13, pop-1, sptf-3*, and *lin-39*, each encoding a transcription factor [40]. The *him-8* mutations suppress only those transcription factor mutations that affect the DNA binding domains, prompting the hypothesis that HIM-8 also has a chromatin remodeling function that affects gene expression [40].

If the interaction of *him-8* with *spe-47* and *spe-50* is due to the chromatin remodeling role for HIM-8 protein, then it might be that the *him-8* mutation is altering the expression of other genes, one or more of which has a more direct interaction with *spe-47* and *spe-50.* We could not identify a sperm defect for the *spe-47; spe-50; him-8* triple mutants other than there were more sperm in the reproductive tract than fertilized eggs produced, suggesting a defect in fusion with the oocytes. SPE-47 localizes to the fibrous body-membranous organelle complexes [FB-MOs; 18], and by its colocalization with SPE-47, so does SPE-50. These complexes are involved in many aspects of sperm development, from acting as vehicles for MSP transport during the meiotic divisions to the remodeling of the spermatid during its transformation to an active spermatozoon. Many gene products are involved in FB-MOs: at least nine different genes have mutant phenotypes that affect FB-MO morphogenesis or function [10]. Here, our results suggest that, in combination with altered gene expression from a HIM-8 deficit, the FB-MO-associated SPE-47 and SPE-50 proteins are important to the ability of sperm to fuse with passing oocytes, even though the two proteins disappear before the spermatids form.

## Acknowledgements

We are grateful to Steven L’Hernault for suggesting the complementation tests in identifying the *spe-50* gene and in providing the *spe-50* designation.

